# Multi-site sampling and risk prioritization reveals the public health relevance of antibiotic resistance genes found in wastewater environments

**DOI:** 10.1101/562496

**Authors:** Chengzhen L. Dai, Claire Duvallet, An Ni Zhang, Mariana G. Matus, Newsha Ghaeli, Shinkyu Park, Noriko Endo, Siavash Isazadeh, Kazi Jamil, Carlo Ratti, Eric J. Alm

## Abstract

The spread of bacterial antibiotic resistance across human and environmental habitats is a global public health challenge. Wastewater has been implicated as a major source of antibiotic resistance in the environment, as it carries resistant bacteria and resistance genes from humans into natural ecosystems. However, different wastewater environments and antibiotic resistance genes in wastewater do not all present the same level of risk to human health. In this study, we investigate the public health relevance of antibiotic resistance found in wastewater by combining metagenomic sequencing with risk prioritization of resistance genes, analyzing samples across urban sewage system environments in multiple countries. We find that many of the resistance genes commonly found in wastewater are not readily present in humans. Ranking antibiotic resistance genes based on their potential pathogenicity and mobility reveals that most of the resistance genes in wastewater are not clinically relevant. Additionally, we show that residential wastewater resistomes pose greater risk to human health than those in wastewater treatment plant samples, and that residential wastewater can be as risky as hospital effluent. Across countries, differences in antibiotic resistance in residential wastewater can, in some cases, reflect differences in antibiotic drug consumption. Finally, we find that the flow of antibiotic resistance genes is influenced by geographical distance and environmental selection. Taken together, we demonstrate how different analytical approaches can provide greater insights into the public health relevance of antibiotic resistance in wastewater.

## 1. Introduction

The spread of antibiotic resistant bacteria and resistance genes is a global public health challenge (The Review on Antimicrobial Resistance, 2014; World Health Organization, 2014; Zaman et al., 2017). Antibiotic resistance genes (ARGs) confer resistance to antibiotics and are present in both bacterial pathogens and the broader environment (Allen et al., 2010; Baquero et al., 2008; D’Costa et al., 2011; Huijbers et al., 2015; Lee et al., 2018). The transfer of these genes between bacteria through horizontal gene transfer is of particular concern, as it can facilitate the transmission of ARGs from environmental reservoirs to human pathogens (Allen et al., 2010; Cantón, 2009; Forsberg et al., 2012; Martínez, 2018; Pehrsson et al., 2016). While current efforts to monitor antibiotic resistance are largely limited to clinical settings (“EARS-Net,” 2010, p., “GLASS,” 2015, “NARMS,” 2018), government agencies have begun to recognize the importance of antibiotic resistance in the environment (Gaze and Depledge, 2017; Wellcome Trust et al., 2018). As health officials explore the need for environmental surveillance of antibiotic resistance, it is critical to understand the human health relevance of ARGs found in different environments.

Efforts to monitor antibiotic resistance in the environment face the challenge of interpreting the threat that environmental resistomes pose to human health (Huijbers et al., 2015). Environmental bacteria are difficult and costly to culture (Larsson et al., 2018; Wellcome Trust et al., 2018). As such, the ability to identify the evolution, transmission, and host range of resistance genes is limited (Larsson et al., 2018). To overcome this challenge, studies have employed metagenomic sequencing to measure antibiotic resistance in the environment (Martínez et al., 2015; Ng et al., 2017; Rowe et al., 2017). These studies often establish the presence of resistance genes in an environment as a risk to human health. Antibiotic resistance genes, however, do not all pose the same risk, and the presence of certain genes may be more indicative of the organisms harboring these genes than of a real public health threat (Martínez et al., 2015). In order for resistance genes to be of relevance to human health, they should reside on mobile genetic elements and be hosted by human bacterial pathogens (Bengtsson-Palme and Larsson, 2015; Martínez et al., 2015). Most resistance genes in the environment, however, are unlikely to be found in human-associated bacteria, as they often occupy different habitats and are phylogenetically distant (Bengtsson-Palme and Larsson, 2015; Larsson et al., 2018; Martínez et al., 2015). Therefore, studies need to move beyond reporting the presence and abundance of ARGs in an environment and also assess their relevance to human health (Bengtsson-Palme and Larsson, 2015; Huijbers et al., 2015; Martínez et al., 2015; Topp et al., 2018).

Wastewater has been proposed as an important source of environmental antibiotic resistance (Marti et al., 2013; Munir et al., 2011; Rizzo et al., 2013; Rodriguez-Mozaz et al., 2015), as well as an environment for monitoring the prevalence of antibiotic resistance in humans (Fahrenfeld and J. Bisceglia, 2016; Gaze and Depledge, 2017; “COMPARE,” 2016). Previous studies of antibiotic resistance in wastewater have primarily focused on hospital effluents and wastewater treatment plants (WWTPs), locations which are widely viewed as hotspots for antibiotic resistant bacteria and horizontal gene transfer (Harris et al., 2014; Ng et al., 2017; Rizzo et al., 2013; Rodriguez-Mozaz et al., 2015; Varela et al., 2014). WWTPs serve as the interface between human society and the environment, as wastewater from various sources meet at WWTPs and undergo treatment processes before being released into the environment. Despite the efforts to remove human-associated bacteria, ARGs are still prevalent in WWTP effluents (Ju et al., 2018; Rizzo et al., 2013; Rodriguez-Mozaz et al., 2015). Hospital wastewater has also been proposed as an important source of environmental antibiotic resistance due to the high levels of antibiotic use and resistant bacteria amongst hospital patients (Harris et al., 2014; Wellcome Trust et al., 2018). Abundances of ARGs in hospital effluents have been found to be higher than in downstream environments such as WWTPs and surface water (Buelow et al., 2018; Ng et al., 2017; Rodriguez-Mozaz et al., 2015; Rowe et al., 2017). However, hospitals contribute less than 1% of the total amount of municipal wastewater at WWTPs and have been found to have little influence on the levels of antibiotic resistance observed at WWTPs (Buelow et al., 2018; Harris et al., 2014; Kümmerer, 2004; Varela et al., 2014). Thus, it remains unclear which wastewater environments should be the focus for monitoring environmental ARGs.

In this study, we combine multi-site sampling of upstream and downstream wastewater with risk prioritization of ARGs to evaluate the public health relevance of antibiotic resistance found in wastewater environments. We show that many ARGs found in wastewater are not human-associated and most genes in wastewater are likely not of risk to human health. We also find that the abundance of ARGs in wastewater for certain classes of antibiotics mirrors antibiotic consumption across countries. Lastly, by comparing resistomes sampled within and across countries, we show that the diversity of resistance genes is shaped by environmental selection and geography. Taken together, this study provides evidence for the hypothesis that different genes and environments pose different risks to human health and illustrates the complexities in interpreting the public health relevance of resistomes.

## 3. Results and Discussion

### 3.1 Residential wastewater resistomes are a complex mixture of human-associated and environmental antibiotic resistance genes

To understand patterns of antibiotic resistance across human-associated environments, we compared the metagenomes of residential wastewater with those from similar wastewater environments and the human gut microbiome. We collected and sequenced wastewater from 13 residential manholes in and surrounding the urban areas of Boston, MA; Seoul, South Korea; and Kuwait City, Kuwait (Table S1; Materials and Methods). We also collected untreated wastewater from the influent of WWTPs in the US and Kuwait. To compare residential wastewater with other wastewater environments and human feces, we downloaded and reprocessed publicly available metagenomic data of adult fecal samples in the US and South Korea and a hospital effluent in the U.K. (Materials and Methods). We used ShortBRED with the SARG database to quantify the abundance of antibiotic resistance genes (ARGs) from metagenomic data (Material and Methods).

Residential wastewater contains more ARGs than human feces and WWTP influent but less than hospital effluent. The median number of ARGs in residential wastewater was 161 (Figure 1A). In comparison, hospital effluent and WWTP influent samples had median ARG counts of 238.5 and 125.5 genes, respectively (Figure 1A). All types of wastewater samples contained more ARGs than fecal samples, which had medians of 50 genes in South Korea and 29.5 genes in the US. We found similar trends in abundance measurements (Figure 1B). Previous studies have implicated both hospital effluent and WWTPs as hotspots for antibiotic resistance, with hospital effluent harboring more ARGs than WWTPs (Ng et al., 2017; Rizzo et al., 2013; Rowe et al., 2017). Our results are consistent with these expectations. The higher levels of antibiotic resistance in residential wastewater than at WWTPs suggest that residential wastewater is also a major reservoir of resistance genes and is a source of antibiotic resistance for WWTPs.

**Figure 1.**
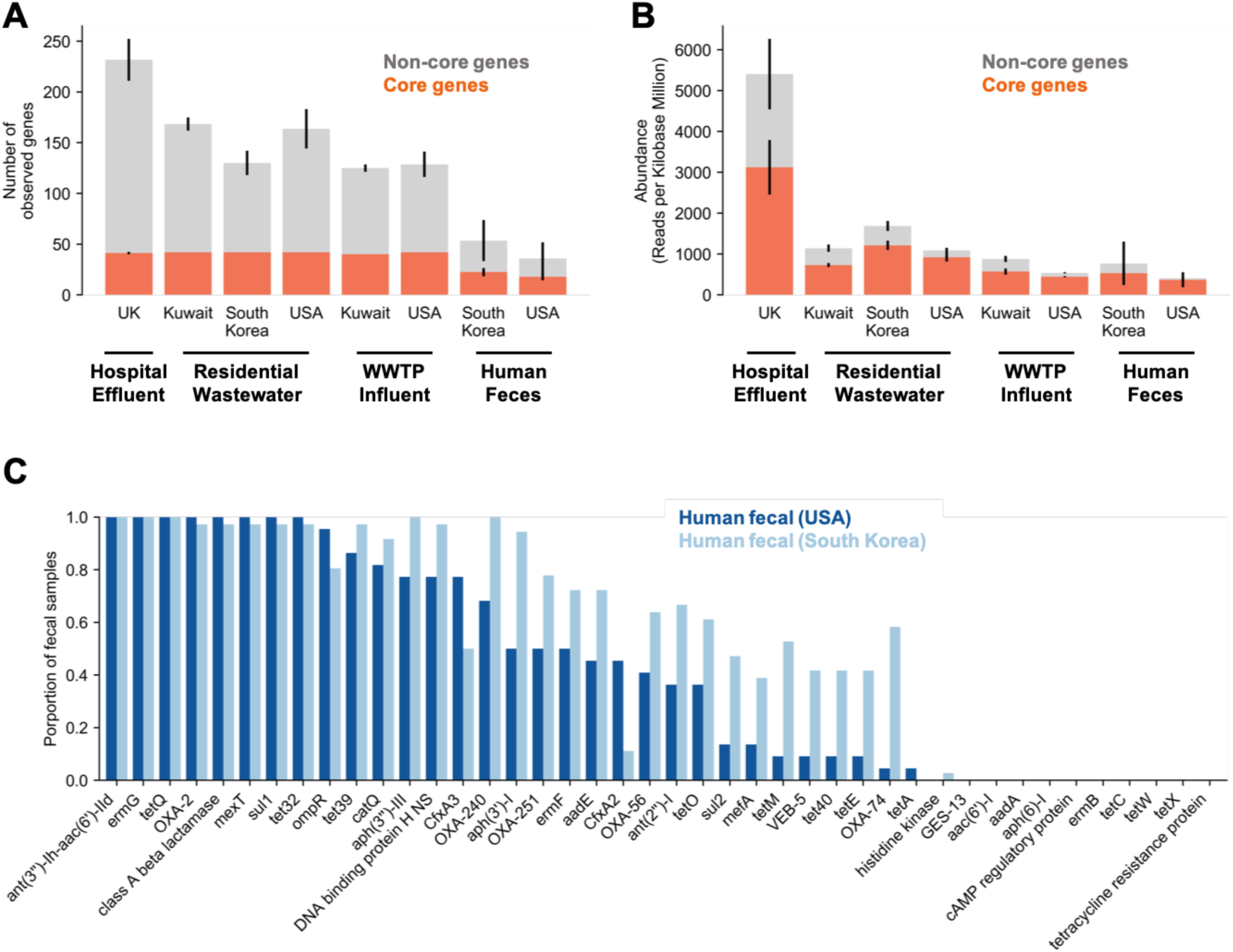
Presence and abundance of antibiotic resistance genes across environments. (A) Number of antibiotic resistance genes observed per environment. Orange coloring represent the core wastewater genes (n = 42) while grey colorings represent the remaining non-core genes. Error bars for each coloring (orange and grey) represent the standard deviations in the number of genes present for that category. (B) Abundance of antibiotic resistance genes observed per environment in reads per kilobase millions. Coloring and error bars represent the same as in (A). (C) Proportion of the human fecal samples in South Korea (n = 36) and the US (n = 22) with each core wastewater gene.

To identify common characteristics of antibiotic resistance in residential wastewater, we looked for genes shared across catchment sites and found a set of 42 genes present in at least one sample from each residential manhole (Figure 1A). These 42 ‘core genes’ accounted for the majority of ARG abundance in residential wastewater (Figure 1B). All of these core genes were also observed at high abundances in hospital effluent and WWTP influent (Figure 1A, 1B). In human fecal samples, these core wastewater ARGs made up approximately 90% of the total ARG abundance (median = 96.2% and 90.8% in US and South Korean individuals, respectively), confirming that the majority of ARGs found in the human gut microbiome are also found in wastewater (Figure 1B). Three South Korean fecal samples had high abundance of ARGs (>50%) that were not part of the core set, with many of these genes conferring multidrug or beta-lactam resistance (Figure S1).

However, not all of the core wastewater genes are found in human feces, suggesting that many of the ARGs measured in wastewater may not be directly relevant to human health. Individual fecal samples only carried approximately half of the 42 core wastewater genes at a time (median = 22 and 18 genes for South Korean and US samples, respectively; Figure 1C). Few core wastewater genes were ubiquitously present in human feces, as only a fifth to a third of the core wastewater genes were found in more than 90% of individuals (14 genes in more than 90% of South Korean fecal samples; 9 in US feces). In fact, 10 of the 42 core genes were not observed in any human fecal sample, suggesting that they are of environmental origin (Figure 1C). Since the transfer of genes from environmental bacteria to human pathogens is less frequent than between human-associated bacteria (Bengtsson-Palme and Larsson, 2015), these 10 core genes likely do not pose an immediate threat to human health. Therefore, not all ARGs identified in wastewater are likely to have equal relevance to human health. Some may reflect genes carried by most humans while others are likely to be derived from the environment.

### 3.2 Most ARGs in wastewater are not clinically-relevant, and upstream wastewater captures more human-related resistomes

To better understand how the presence of ARGs in residential wastewater relates to human health risks, we categorized genes based on their potential pathogenicity. We used the method described in Zhang et al. (in preparation), which ranks the risk of gene variants based on the variant’s observed host pathogenicity, mobility, host range, and anthropogenic prevalence (Materials and Methods). Variants classified into the top two ranks with this method are variants which have been previously observed on plasmids and in a phylogenetically diverse range of hosts, including human pathogens. We therefore refer to the variants in these two ranks as clinically-relevant, as there exists published evidence of these mobile genes posing a substantial risk for the dissemination of resistance (Martínez et al., 2015).

Most of the ARGs found in wastewater and humans are not clinically-relevant, and the majority of the core wastewater genes are also not clinically-relevant. We found that clinically-relevant variants make up less than 50% of the total antibiotic resistance gene abundance in all of the wastewater and human samples we surveyed (Figure 2, S2). These results also held when we looked at the top three ranks, which represents all mobile ARGs (Figure S3). Among the 42 core wastewater genes, clinically-relevant variants were found for only 11 of the core wastewater genes and represented less than 15% of the total ARG abundance in residential wastewater (Figure 2, light red). These results directly support the hypothesis proposed by Martínez et al. (2015), in which the number of antibiotic resistance genes that are actually acquired by human pathogens and lead to clinical complications are low compared to the number of sequences classified as resistance genes in metagenomic studies. Thus, simply quantifying the presence and abundance ARGs in a given sample does not necessarily measure the health relevance of that sample’s resistome.

**Figure 2.**
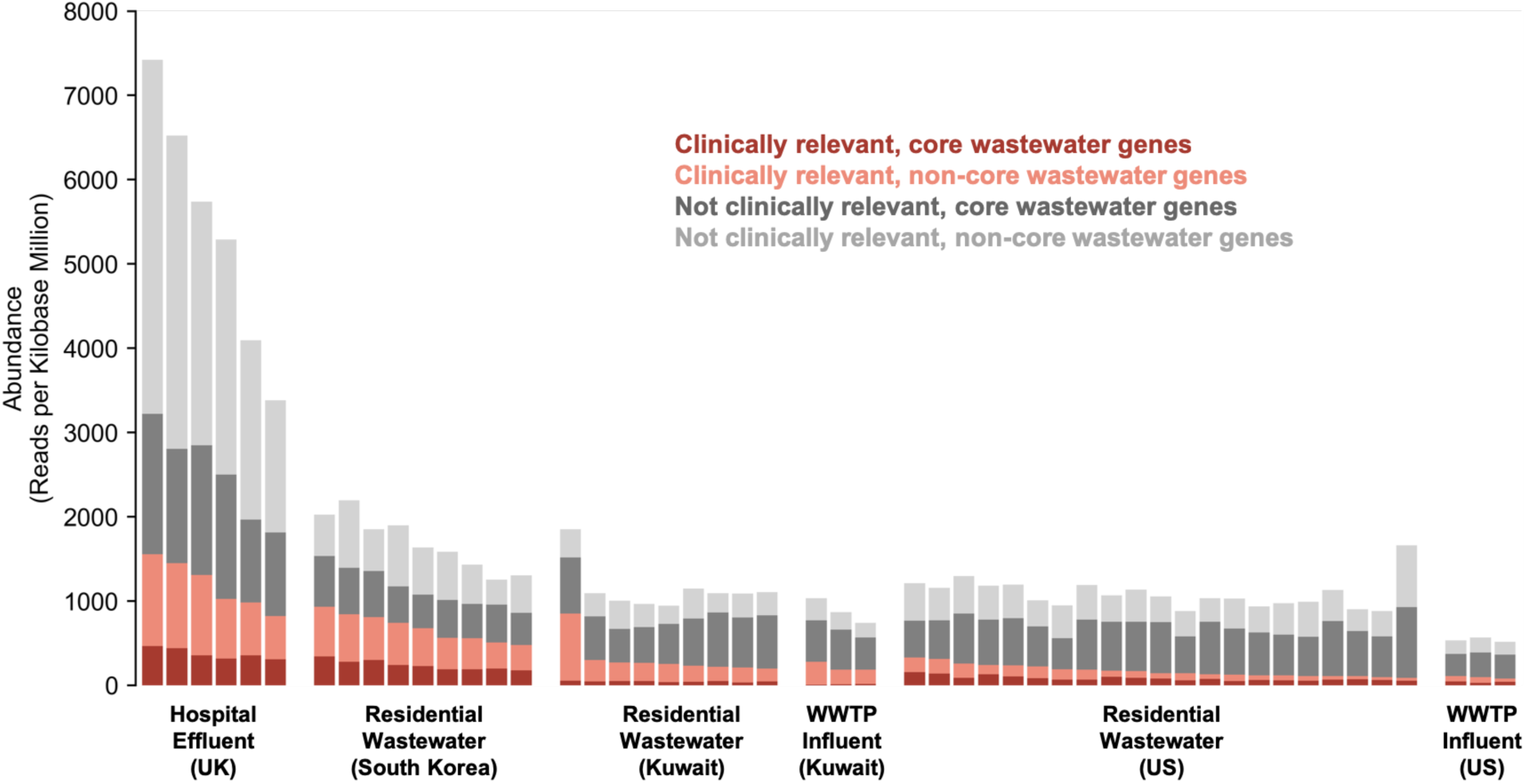
Risk prioritization of antibiotic resistance genes in wastewater environments. Abundance of antibiotic resistance gene in each wastewater environment, categorized by risk. Proportions in shades of red represent abundances of clinically-relevant antibiotic resistance genes while shades of grey represent non-clinically-relevant genes. Within each coloring (red and grey), genes in the core set of wastewater genes are shaded darker (i.e., dark red = genes which are clinically relevant and core; dark grey = genes which are not clinically relevant and core). Samples within each environment are ranked in descending order based on total abundance of clinically relevant genes. Genes are defined as clinically-relevant if they have previously been observed on plasmids and have a phylogenetically diverse range of hosts. Abundances are measured in reads per kilobase million.

All types of upstream wastewater harbor more clinically-relevant ARGs than the influent of WWTPs. We compared the presence and abundance of clinically-relevant variants in residential wastewater to the traditionally studied environment of wastewater treatment plants. In both countries (Kuwait and US) where we sampled upstream and downstream wastewater, residential wastewater had higher abundances of clinically-relevant variants than the respective WWTP influent (Figure 2). At the same time, WWTP influent contained fewer variants found in human feces than residential wastewater. We focused on the US where data was available for human fecal, residential wastewater, and WWTP influent samples. We found that of the 29 clinically-relevant variants present in at least one healthy fecal sample, 90% (n = 26) were present in residential wastewater. In contrast, only 76% (n = 22) were found in WWTP influents. More generally, 82% (n = 270) of all of the variants found in fecal samples (n = 328) were observed in residential wastewater, whereas only 59% (n = 195) of them were identified downstream in the WWTP influents. The microbial composition of human waste is known to decrease in similarity to the human fecal microbiota as it progresses through the sewage system (Pehrsson et al., 2016). Our results suggest this decrease also occurs for resistomes, both clinically-relevant and non-clinically-relevant ones, as upstream sampling better reflects the collection of microbes and antibiotic resistance in the contributing human population.

Although hospital effluent has more overall ARGs and is traditionally thought of as a unique hotspot for environmental antibiotic resistance, we found that residential wastewater can harbor as many clinically-relevant variants as hospital effluent. U.K. hospital effluent had abundances of clinically-relevant variants ranging from 821 to 1555 RPKM (Figure 2). By comparison, the residential wastewater of South Korea and Kuwait had abundances of clinically-relevant variants ranging from 478 to 933 RPKM and 200 to 850 RPKM, respectively (Figure 2). Previous studies implicating hospital wastewater as a major source of environmental antibiotic resistance have largely compared hospital wastewater with environments further downstream such as wastewater treatment plants or surface water (Buelow et al., 2018; Ng et al., 2017; Rodriguez-Mozaz et al., 2015; Rowe et al., 2017). Our results thus challenge the prevailing hypothesis that hospital effluent represents the most concerning source of environmental antibiotic resistance genes and suggests that all upstream wastewater may serve as important reservoirs of clinically-relevant genes.

### 3.3 Antibiotic resistance patterns across countries can reflect human activity

To assess whether ARGs in residential wastewater reflect population-level antibiotic consumption, we compared the abundance of antibiotic resistance genes with available consumption data from the IQVIA MIDAS database (Klein et al., 2018). We limited our analysis to South Korea and the US, where consumption data included both hospital and non-hospital use.

For certain antibiotic classes, resistance across countries reflects antibiotic consumption patterns. Consumption of aminoglycoside and beta-lactam antibiotics is higher in South Korea than the US, as reflected by the median abundance of resistance genes to these antibiotics in the respective residential wastewater and fecal samples (Figure 3A, top and bottom row). South Korean samples also had higher abundances of chloramphenicol resistance than US samples. While differences in current antibiotic consumption is negligible, historical consumption of chloramphenicol was higher in South Korea. Unlike the US, which phased out chloramphenicol use in the 1960s, South Korea continued its usage until 2013. The presence of chloramphenicol resistance may therefore be the result of persistent resistance, as previous studies have found that resistance genes can persist for a long time after their introduction into the microbial flora (Forslund et al., 2013). However, this hypothesis needs further validation with better consumption data and direct antibiotic susceptibility testing. These resistance patterns were also observed in the human fecal samples (Figure 3A, middle row) and were more evident in clinically-relevant genes than non-clinically-relevant ones (Figure S4).

**Figure 3.**
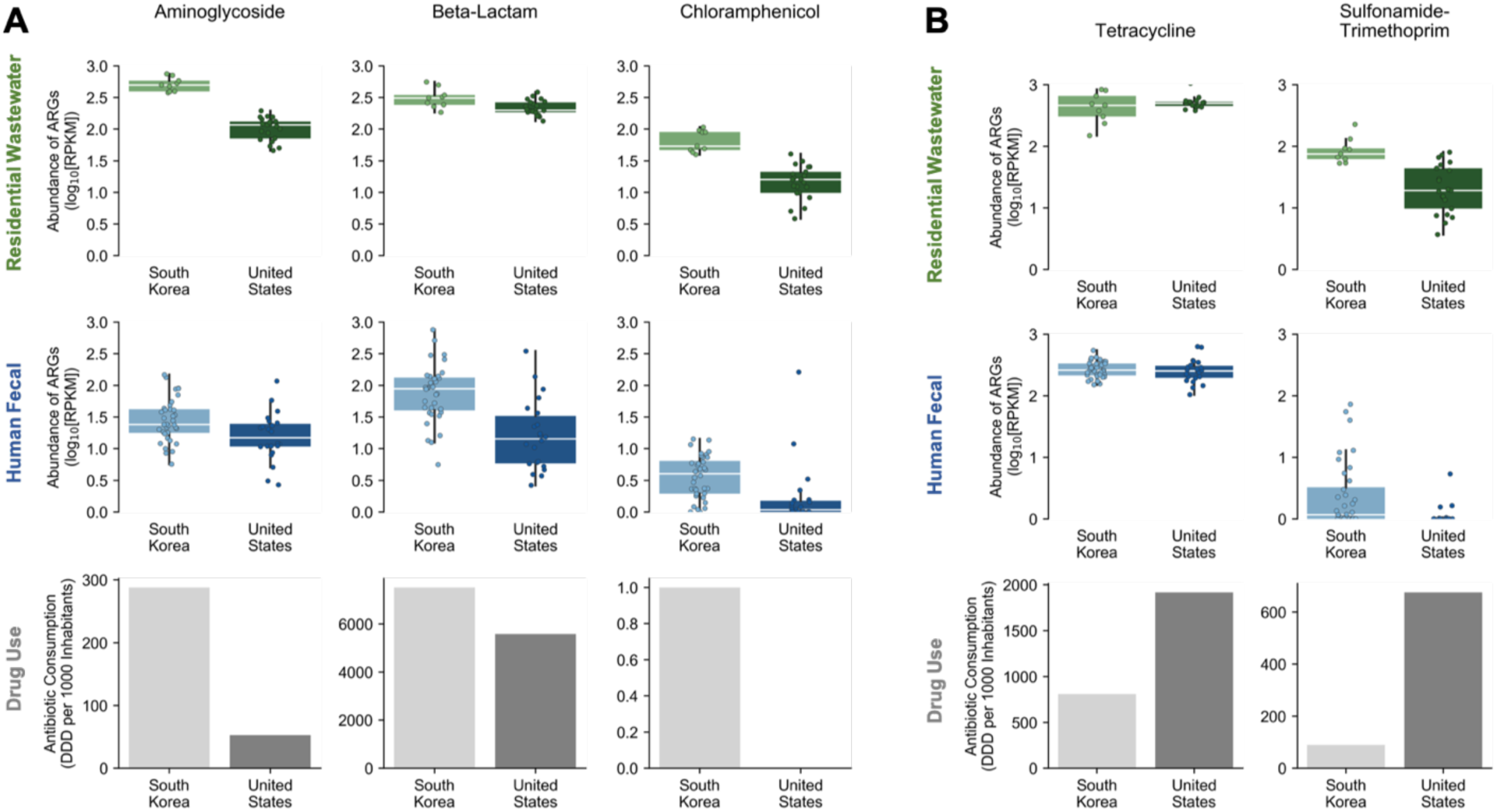
Antibiotic resistance and antibiotic consumption across geography by drug class. Patterns of antibiotic resistance in residential wastewater (top panels) and human feces (middle panels) and patterns antibiotic consumption (bottom panels) between South Korea and the United States. Antibiotic resistance and antibiotic consumption pattern show concordance for some classes of antibiotics (A) but not for others (B). Antibiotic resistance is represented in log abundances of reads per kilobase million (RPKM) while antibiotic consumption is represented in defined daily dose (DDD) per 1,000 inhabitants. Each point represents one sample. The y-axis range for each antibiotic consumption plot is different. Data for antibiotic consumption (2015) was obtained from the IQVIA MIDAS database (Klein et al., 2018; https://resistancemap.cddep.org/AntibioticUse.php).

Other antibiotic classes, however, have resistance patterns which do not reflect known country-level differences in antibiotic consumption. Sulfonamide-trimethoprim consumption is higher in the US than South Korea, but resistance to sulfonamide and trimethoprim, both separately and combined, were higher in South Korean samples (Figure 3B). Similarly, tetracycline consumption is higher in the US than South Korea but median abundance of tetracycline resistance genes showed the opposite trend. These inconsistencies emphasize how multiple factors contribute to the antibiotic resistance observed in the environment (Collignon et al., 2018). Other factors, such as antibiotic use in agriculture and environmental contamination with antibiotics, can also drive resistance but data on them is limited (Berendonk et al., 2015; Hoelzer et al., 2017). As efforts are made to fully understand the public health relevance of environmental antibiotic resistance, more comprehensive data on antibiotic use across sectors as well as better approaches to measuring antibiotic resistance in the environment are needed.

### 3.4 Flow of antibiotic resistance genes varies across geographic scales

The composition of resistomes differs between residential catchment sites within a city. Pairs of samples from different residential manholes in the same city had higher beta diversity than sample pairs from the same manhole (median JSD = 0.30 vs. 0.27, respectively; p < 0.01, PERMANOVA; Figure 4A). Samples from the same manhole were collected one after another with ∼5 minute breaks in between (Materials and Methods). Therefore, different sets of humans likely contributed to the ARGs in each sample, resulting in the observed differences between these samples (Ort et al., 2005). Across manholes, however, the physical conditions of the wastewater environment are also different, with levels of oxygen and temperature often varying between sites (Wellcome Trust et al., 2018). These variations likely contribute to differences in microbial composition and consequently, ARG composition (Huisman et al., 2004; Pehrsson et al., 2016; Vollertsen and Hvitved-Jacobsen, 1999). As expected, beta diversity between samples from different countries are higher than all within-city comparisons (median JSD = 0.53; P < 0.01, PERMANOVA; Figure 4A). Thus, these differences in resistome composition may reflect selection resulting from different environmental conditions in individual manholes.

**Figure 4.**
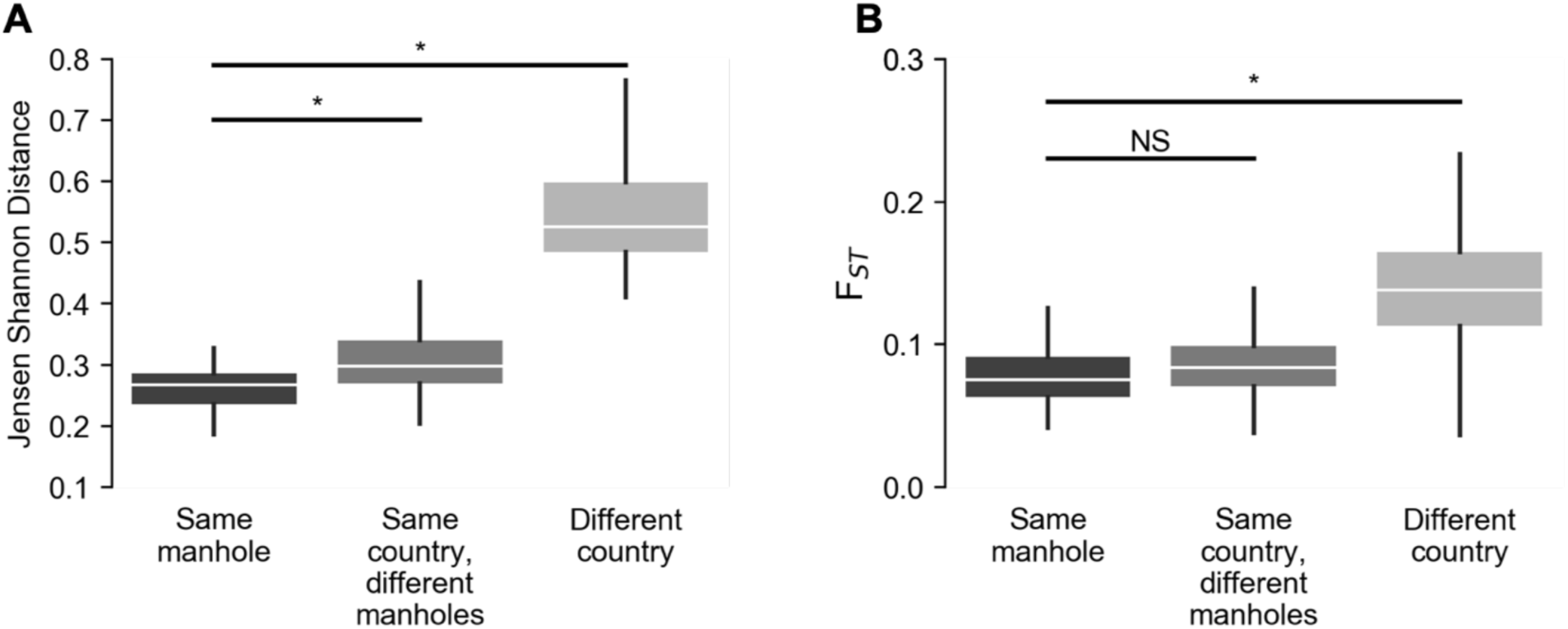
Beta diversity and nucleotide diversity of antibiotic resistance genes across different geographical scales. (A) Beta diversity in Jensen Shannon Distance of resistomes between samples at different scales of geographical comparisons. *P < 0.01 (B) Nucleotide diversity of resistomes between samples at different scales of geographical comparisons. Nucleotide diversity between each pair of samples was measured in terms of the average F_ST_ of genes with sufficient coverage and polymorphism (n = 35; Material and Methods). *P < 0.01. NS = not significant.

Despite differences in the overall resistome composition between different catchments, the nucleotide diversity of individual ARGs remain similar across manholes in the same city. We aligned metagenomic reads against representative nucleotide sequences to identify single nucleotide polymorphisms for each antibiotic resistance gene (Materials and Method). We then calculated F_ST_ values to quantify genetic diversity between samples for each gene that had sufficient coverage and polymorphisms (N = 35; Materials and Method). F_ST_ is a measure of genetic differentiation, with values ranging from 0 to 1 where 0 represents no substructure and 1 means completely different alleles between the subpopulations (Wright, 1951). Overall, pairs of samples from different manholes within the same city did not have significantly different F_ST_ values than pairs of samples from the same manhole (median F_ST_ = 0.075 versus 0.083, respectively; P = 0.06, PERMANOVA). That is, variants of a gene present across multiple locations within a city were as similar to each other as those found in consecutive samples taken from one manhole. As expected, genes were more dissimilar across countries, suggesting that there exists barriers to the distribution of ARGs across larger geographic distances (median F_ST_ = 0.14 across country versus median F_ST_ = 0.08 within country; P < 0.01, PERMANOVA; Figure 4B). To understand how the composition of these genes differ across catchment sites and across geography, we evaluated the beta diversity of these 35 genes between samples. Similar to the results for the overall resistome composition, beta diversity was highest between countries and was higher between different catchments in the same city than within the same catchment (Figure S5). Taken together, these results suggest that while individual manhole environments play a role in selecting for abundances of genes, specific gene variants themselves likely have few barriers to distribution at smaller geographical scales.

## 4. Conclusion

Urban sewage systems play a major role in the dissemination of antibiotic resistance from humans to the environment. In this study, we evaluated the presence of antibiotic resistance genes across wastewater environments, assessing their relevance and risk to human health and identifying patterns across geography. We sampled upstream residential wastewater, an understudied part of the sewage system that is close to the human waste sources, across multiple countries and compared it with other wastewater environments to highlight challenges in the evaluation of antibiotic resistance in wastewater. We found that a substantial proportion of the antibiotic resistance genes commonly found in wastewater are not present in human feces and do not pose an immediate threat to human health, suggesting that evaluating an environment’s risk to human health should not rely solely on measuring the presence of resistance genes. While WWTPs and hospital effluents are traditionally viewed as antibiotic resistance hotspots, we showed that residential wastewater may also be a major source of environmental resistance, containing higher abundances of ARGs than WWTP samples and at times reaching comparable levels of risk as hospital effluent. Although some classes of antibiotics exhibited similar patterns between consumption and resistance across countries, others did not, highlighting that the relationship between environmental antibiotic resistance and population-level human antibiotic consumption is complex. Finally, we demonstrated that despite some differences due to environmental selection between manholes, gene flow readily occurs within a city but larger geographical distances serve as a barrier to gene flow.

Overall, our study highlights the challenges in evaluating the public health relevance of antibiotic resistance genes found in wastewater environments and provides preliminary insights on how to address these challenges. Targeting specific genes (e.g., human-associated genes, clinically-relevant genes) and evaluating resistomes at different scales (e.g., nucleotide diversity, compositional differences) reveals relationships between antibiotic resistance found in the environment and human health. As wastewater becomes a focal point in efforts to monitor population-level antibiotic resistance, targeted approaches such as those presented here can be incorporated into surveillance efforts to yield more actionable public health insights.

## 2. Material and Methods

### 2.1 Wastewater collection and processing

Grab wastewater samples (250 mL) were taken from the manholes at 13 different residential catchment sites in Boston, MA (3 neighborhoods); Cambridge, MA (4 neighborhoods); Seoul, South Korea (3 neighborhoods); and the surrounding areas of Kuwait City, Kuwait (3 neighborhoods). We also collected raw wastewater samples at a pump station directly feeding into a WWTP in Chelsea, MA and at a WWTP in Sulaibiya, Kuwait (Table S1). The pump station in Chelsea serves parts of the Boston metropolitan area while the Sulaibiya WWTP serves parts of Kuwait City and its surrounding area. We consider samples from these two locations as WWTP influent samples as they are untreated. WWTP influent samples were collected using a sampling pole while samples from manholes were collected using a commercial peristaltic pump (Boxer) at a sampling rate of 50 mL/min. 30 ml of collected wastewater were filtered through 0.2-m PTFE membrane filters (Millipore). PTFE membrane filters were kept in RNAlater at - 80 degrees Celsius until DNA extraction. The lab filtration system consisted of a Masterflex peristaltic pump (Pall), Masterflex PharMed BPT Tubing (Cole-Palmer), 47 mm PFA filter holders (Cole-Palmer) and 47mm PTFE Omnipore filter membranes (Millipore).

### 2.2 DNA extraction and shotgun metagenomic sequencing

0.2-m filter membranes were thawed on ice. RNAlater was removed and filters were washed with phosphate-buffered saline (PBS) buffer twice. Metagenomic DNA was extracted from each filter using the Power Water extraction kit (MO BIO Laboratories Inc.), according to manufacturer’s instructions. Sample concentrations were quantified with the Quant-iT PicoGreen dsDNA Assay (Life Technologies) and normalized to equal concentration. Paired-end libraries were prepared with 100-250 pg of DNA using the Illumina Nextera XT DNA Library Preparation Kit according to the manufacturer’s instructions. Insert sizes and concentrations for each library were determined with an Agilent Bioanalyzer DNA 100 kit (Agilent Technologies). Libraries were finally sequenced on the Illumina NextSeq platform at the MIT Biomicro Center to generate 2 × 150 bp paired reads. Approximately 10 million reads were generated for each sample (∼5 million for each pair).

### 2.3 Shotgun metagenomic sequencing data processing

Raw paired-end DNA sequences (FASTQ reads) were quality controlled prior to any analysis. Low-quality reads and adaptor sequences were removed using Trimmomatic version 0.36 with the ILLUMINACLIP parameters: NexteraPE-PE.fa:2:30:10 SLIDINGWINDOW:4:20 MINLEN:50 (Bolger et al., 2014). To remove sequences resulting from human contamination, we used Bowtie 2 version 2.3.0 in default mode (Langmead and Salzberg, 2012). We aligned trimmed reads to the human reference genome (GRCh38) and removed the reads that mapped from downstream analyses.

### 2.4 Comparison of wastewater resistomes to published fecal and wastewater samples

For comparison with hospital effluent wastewater and human feces, whole metagenome shotgun reads were downloaded from ENA or SRA. Hospital effluent wastewater sequences was obtained from Rowe et al. (2017), ENA accession PRJEB12083; Human fecal samples were obtained from Lim et al. (2014), ENA accession ERP002391, for South Korea and Obregon-Tito et al. (2015), SRA accession PRJNA268964, for the United States. Raw FASTQ reads from these samples were processed as above and all further analyses were performed together with sequences generated in this study.

### 2.5 Quantification of antibiotic resistance genes

To identify and quantify the abundance of antibiotic resistance genes in each quality-controlled metagenomic sample, we used ShortBRED version 0.9.5 (Kaminski et al., 2015). ShortBRED first clusters proteins based on shared amino acid identity and identifies unique marker sequences for each cluster that distinguish them from close homologues. ShortBRED then quantifies the abundance of each protein cluster by mapping reads to those unique markers. Since antibiotic resistance genes can share homology with genes of non-resistance functions, ShortBRED provides greater accuracy than mapping reads to the entire protein sequence.

We generated ShortBRED markers from the Structured ARG (SARG) reference database (Yang et al., 2016). The SARG database contains 4,049 amino acid sequences and was constructed by integrating the ARDB and CARD databases (Yang et al., 2016). We used 100% identity for clustering and mapped the clustered sequences to the Integrated Microbial Genomes database version 3.5 to generate the set of unique markers (Kaminski et al., 2015). We used the parameters: --id 1.0 --clustid 1.0 --qclustid 1.0. In total, ShortBRED generated 9,807 marker sequences. Finally, these marker sequences were used to quantify antibiotic resistance gene abundance in metagenomes by mapping paired FASTQ reads to them at 99% sequence identity. Abundances were normalized to reads per kilobase per million reads (RPKM).

### 2.6 Risk Ranking of Antibiotic Resistance Genes

We used ARG Ranker to prioritize antibiotic resistance genes based on their risk relevance to human health (Zhang et al., in preparation; https://github.com/caozhichongchong/arg_ranker). Building off of the risk ranking criteria outlined in Martínez et al. (2015), ARG Ranker takes the genes in the SARG database and assesses their prevalence in publicly available whole genome and metagenomic sequences. Since transferability is a major bottleneck that determines an ARG’s risk potential, genes found on the plasmids of previously-sequenced pathogen isolates are ranked higher in risk potential (Rank 1-3) than genes not found in plasmids (Rank 4-5). Amongst these mobile genes (Rank 1-3), those with a phylogenetically diverse range of hosts are ranked higher (Rank 1-2) than those with a less phylogenetically diverse range of hosts (Rank 3). We consider the genes in the top two ranks as clinically-relevant as they have the greatest potential for transfer between host and environments. Abundance in anthropogenic environments versus natural environments differentiate genes between the top two ranks, with genes having higher abundances being ranked higher.

### 2.7 Identification of single nucleotide variants

We used a combination of Bowtie 2 and metaSNV to identify and quantify single nucleotide variants. We first aligned metagenomic reads to a nucleotide database of representative sequence for antibiotic resistance genes using Bowtie 2 version 2.3.0 (Langmead and Salzberg, 2012). This database included a single nucleotide reference sequence for each gene found by ShortBRED. In total, 264 ARGs were assessed. We used the nucleotide sequences from the CARD database that shared > 99% similarity width the SARG database (n = 3136), as the SARG database only included amino acid sequences and was constructed using CARD (Jia et al., 2017). We called and filtered variants with metaSNV version 2.0 using the default setting (Costea et al., 2017).

### 2.8 Statistical analyses

Beta diversity between residential wastewater samples was calculated at gene level using the Jensen-Shannon distance (JSD). Differences in antibiotic resistance gene composition across manholes and countries was evaluated using the PERMANOVA test implemented in the Python package *scikit-bio* v0.4.2 (skbio.stats.distance.permanova).

Nucleotide diversity was measured using Wright’s F_ST_ (Wright, 1951), as implemented in metaSNV v.2.0 with the flag: -div. We only analyze genes present in all three countries (n = 35). Comparisons between manholes and between countries were made using the PERMANOVA test implemented in the Python package *scikit-bio* v0.4.2 (skbio.stats.distance.permanova).

## Code and data availability

All custom scripts generated in Python to analyze the data in this paper are available through GitHub (https://github.com/chengdai/amr_risk). Metagenomics sequencing data generated in this study are available at NCBI Short Read Archive under Bioproject PRJNA524435.

## Supplementary Data

Supplementary data related to this article can be found in the online version.

## Acknowledgements

We thank the Boston Public Works Department, the Cambridge Department of Public Works, the Kuwait Ministry of Public Works, and the Seoul Metropolitan Government for their help in selecting and accessing manholes along our team. We also thank Gwangpyo Ko for allowing us to use his facility in Seoul, South Korea. This work was supported by the Kuwait Foundation for the Advancement of Sciences. C.D. acknowledges support from the Siebel Scholars Foundation. S.I. acknowledges support from the Natural Sciences and Engineering Research Council of Canada. The MIT Center for Microbiome Informatics and Therapeutics supported the computational resources for this project.

## Author contributions

C.L.D., C.D., and E.J.A. designed this study. M.G.M., N.G., S.P. collected the raw samples. N.E. helped select the sites for sampling. M.G.M. processed the samples for sequencing. C.L.D. analyzed the data. C.D., A.Z., and S.I. assisted with data analysis. C.L.D., C.D., and E.J.A. wrote the paper. All authors reviewed and edited the paper. This study was conceived by E.J.A., C.R., and K.J.

